# Predicting cancer outcomes from histology and genomics using convolutional networks

**DOI:** 10.1101/198010

**Authors:** Pooya Mobadersany, Safoora Yousefi, Mohamed Amgad, David A Gutman, Jill S Barnholtz-Sloan, Jose Enrique Velazquez Vega, Daniel J Brat, Lee AD Cooper

## Abstract

Cancer histology reflects underlying molecular processes and disease progression, and contains rich phenotypic information that is predictive of patient outcomes. In this study, we demonstrate a computational approach for learning patient outcomes from digital pathology images using deep learning to combine the power of adaptive machine learning algorithms with traditional survival models. We illustrate how this approach can integrate information from both histology images and genomic biomarkers to predict time-to-event patient outcomes, and demonstrate performance surpassing the current clinical paradigm for predicting the survival of patients diagnosed with glioma. We also provide techniques to visualize the tissue patterns learned by these deep learning survival models, and establish a framework for addressing intratumoral heterogeneity and training data deficits.

## INTRODUCTION

Histology has been an important tool in cancer diagnosis and prognostication for more than a century. Anatomic pathologists evaluate histology for characteristics like nuclear atypia, mitotic activity, cellular density, and tissue architecture, incorporating cytologic details and higher-order patterns to classify and grade lesions. Although prognostication increasingly relies on genomic biomarkers that measure genetic alterations, gene expression, and epigenetic modifications, histology remains an important tool in predicting the future course of a patient’s disease. The phenotypic information present in histology reflects the aggregate effects of molecular alterations on cancer cell behavior, and provides a convenient visual readout of disease aggressiveness. However, human assessments of histology are highly subjective and not repeatable, hence computational analysis of histology imaging has received significant attention. Aided by advances in slide scanning microscopes and computing, a number of image analysis algorithms have been developed for grading (1–4), classification (5–10), and prediction of future metastasis (11) in multiple cancer types.

Deep convolutional neural networks (CNNs) have emerged as an important image analysis tool, and have shattered performance benchmarks in many challenging applications (12). The ability of CNNs to learn predictive features from raw image data is a paradigm shift that presents exciting new opportunities in medical imaging (13–15). Medical image analysis applications have heavily relied on *feature engineering* approaches where algorithms are designed to delineate or detect structures of interest, and to measure pre-defined characteristics of these structures that are believed to be predictive. In contrast, the *feature learning* paradigm of CNNs does not rely on biased a priori definitions of features, and does not require the explicit delineation of anatomic structures which is often confounded by artifacts and variations in image acquisition. While feature learning has become the dominant paradigm in general image analysis tasks, medical applications present unique challenges. Large amounts of labeled data are needed to train CNNs, and medical applications often suffer from data deficits that hurt performance. As “black box” models, CNNs are also difficult or impossible to deconstruct, and so their prediction mechanisms cannot be understood. Despite these challenges, CNNs have been successfully used extensively for medical image classification and segmentation applications (9, 11, 16–24).

Many important problems in the clinical management of cancer involve *time-to-event* prediction, including accurate prediction of overall survival, time to progression, and time to metastasis. Despite overwhelming success in other applications, deep learning has not been widely applied to these problems. Survival analysis has often been approached as a machine-learning classification problem by dichotomizing outcomes (e.g. alive vs. deceased at 5 years) (25). Neural network based Cox regression approaches were explored in early work with low-dimensional clinical datasets, but subsequent analysis found no improvement over basic Cox regression (26). Deeper networks that are capable of feature learning were recently adapted to optimize Cox proportional hazard likelihood and were shown to have superior performance in predicting overall survival from high-dimensional genomic signatures (27), and with low-dimensional clinical datasets (28). For images, similar convolutional networks were applied to predicting overall survival using lung cancer histology but achieved only marginally better than random prediction accuracy (0.629 c-index), and were not compared to simple models based on clinical predictors like age, sex, stage or histologic grade (29).

In this paper, we present a convolutional network based approach called *Survival Convolutional Neural Networks* (SCNN) that can predict overall survival and other time-to-event outcomes from histology images with accuracy that equals or surpasses clinical paradigms based on genomic biomarkers and manual histologic grading. We provide a new training and prediction framework based on image resampling that significantly improves prediction accuracy by mitigating the effects of intratumoral heterogeneity and deficits in the amounts of labeled training data. We also illustrate how genomic and histology imaging data can be integrated into a single SCNN prediction model to significantly improve prognostic accuracy. Finally, we show how the prediction mechanisms of SCNN models can be interpreted using whole-slide risk heatmaps that visualize the risks associated with various regions in a histologic specimen. We systematically validate these approaches by building models to predict overall survival in gliomas using data from The Cancer Genome Atlas Lower Grade Glioma (LGG) and Glioblastoma (GBM) projects.

## RESULTS

### Learning patient outcomes with deep survival convolutional neural networks

The SCNN model architecture is depicted in **Figure 1** (see Figure S1 for detailed diagram). Hematoxylin and eosin tissue sections are first digitized to large whole-slide-images. These images are reviewed using a web-based platform to identify regions-of-interest (ROIs) with representative histologic characteristics (30). High-power fields (HPFs) from these ROIs are then used to train a deep convolutional network that is seamlessly integrated with a Cox proportional hazards model to predict patient outcomes. The network is composed of interconnected layers of image processing operations and nonlinear functions that sequentially transform the HPF image into highly-predictive prognostic features. Convolutional layers first extract visual features from the field image at multiple scales using convolutional kernels and pooling operations. These image-derived features feed into fully-connected layers that perform additional transformations, and then a final Cox model layer outputs a prediction of patient risk. The interconnection - weights and convolutional kernels are trained by comparing risk predicted by the network with survival or other time-to-event outcomes using a backpropagation technique to optimize the statistical likelihood of the network (see Methods).

**Figure 1.**
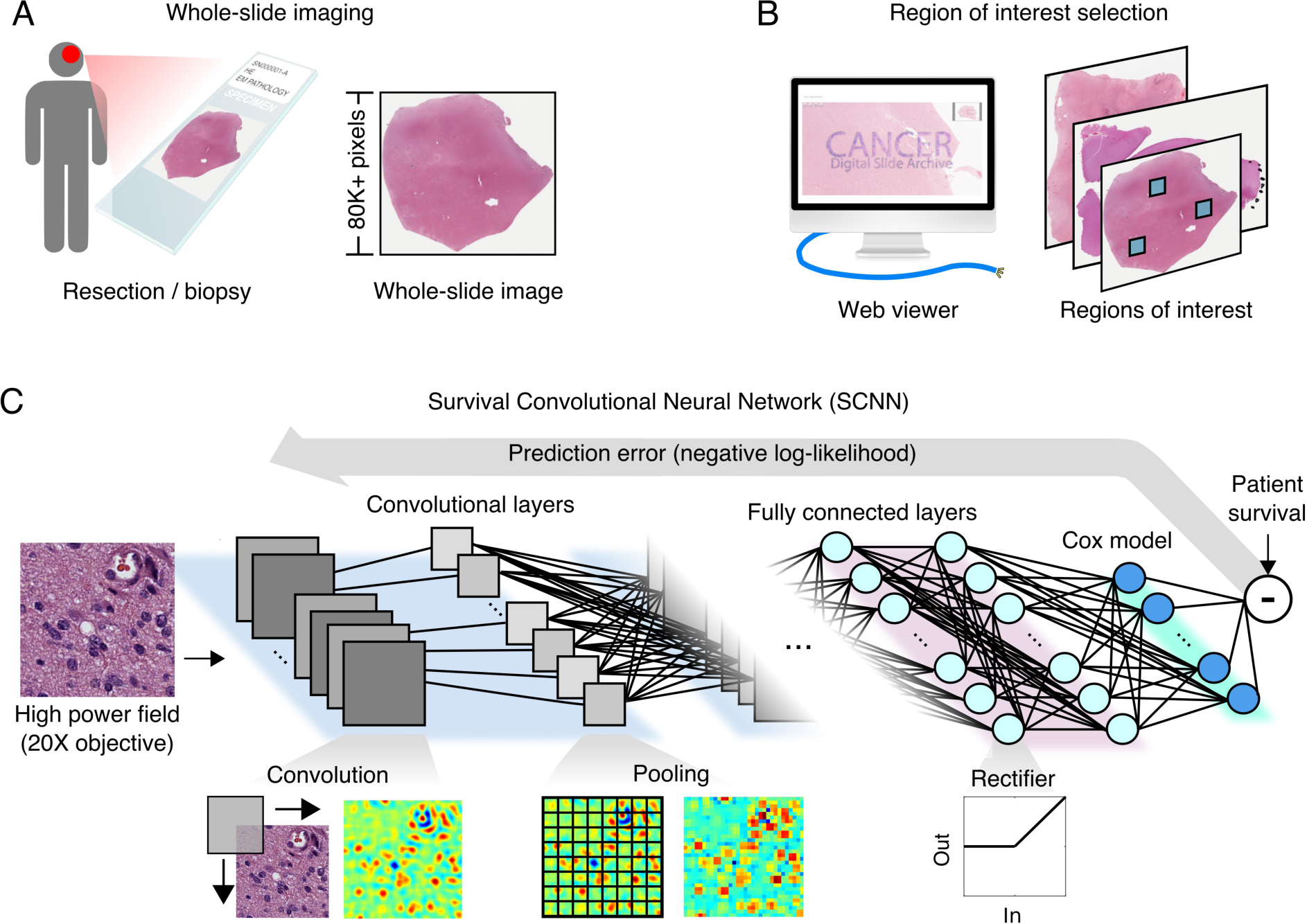
The survival convolutional neural network (SCNN) model. The SCNN combines deep learning convolutional neural networks with traditional survival models to learn survival-related patterns from histology images. **(A)** Large whole-slide images are generated by digitizing hematoxylin & eosin glass slides. **(B)** A web-based viewer is used to manually identify representative regions-of-interest in the image. **(C)** High-power fields are sampled from these regions and used to train a neural network to predict patient survival. The SCNN consists of: 1. Convolutional layers that learn visual patterns related to survival using convolution and pooling operations 2. Fully connected layers that provide additional non-linear transformations of extracted image features and 3. A Cox proportional hazards layer that models time-to-event data like overall survival or time-to-progression. Predictions are compared to patient outcomes to adaptively train the network weights that interconnect each layer.

To improve the performance of SCNN models, we developed resampling techniques to address the limited availability of training samples and intratumoral heterogeneity (see **Figure 2**). For training, new HPFs are randomly sampled from each ROI at the start of each training iteration, providing the SCNN model with a fresh look at each patient’s histology and capturing heterogeneity within the ROI. The SCNN is also trained using multiple such HPFs for each patient (one for each region) to further account for intratumoral heterogeneity across ROIs. For predicting the risk of a new patient with unknown survival, we integrate information from many HPFs by randomly sampling multiple fields in each ROI and using averaging and ranking procedures to create a robust patient level prediction that rejects outlying risk predictions. These resampling procedures are described in detail in Methods.

**Figure 2.**
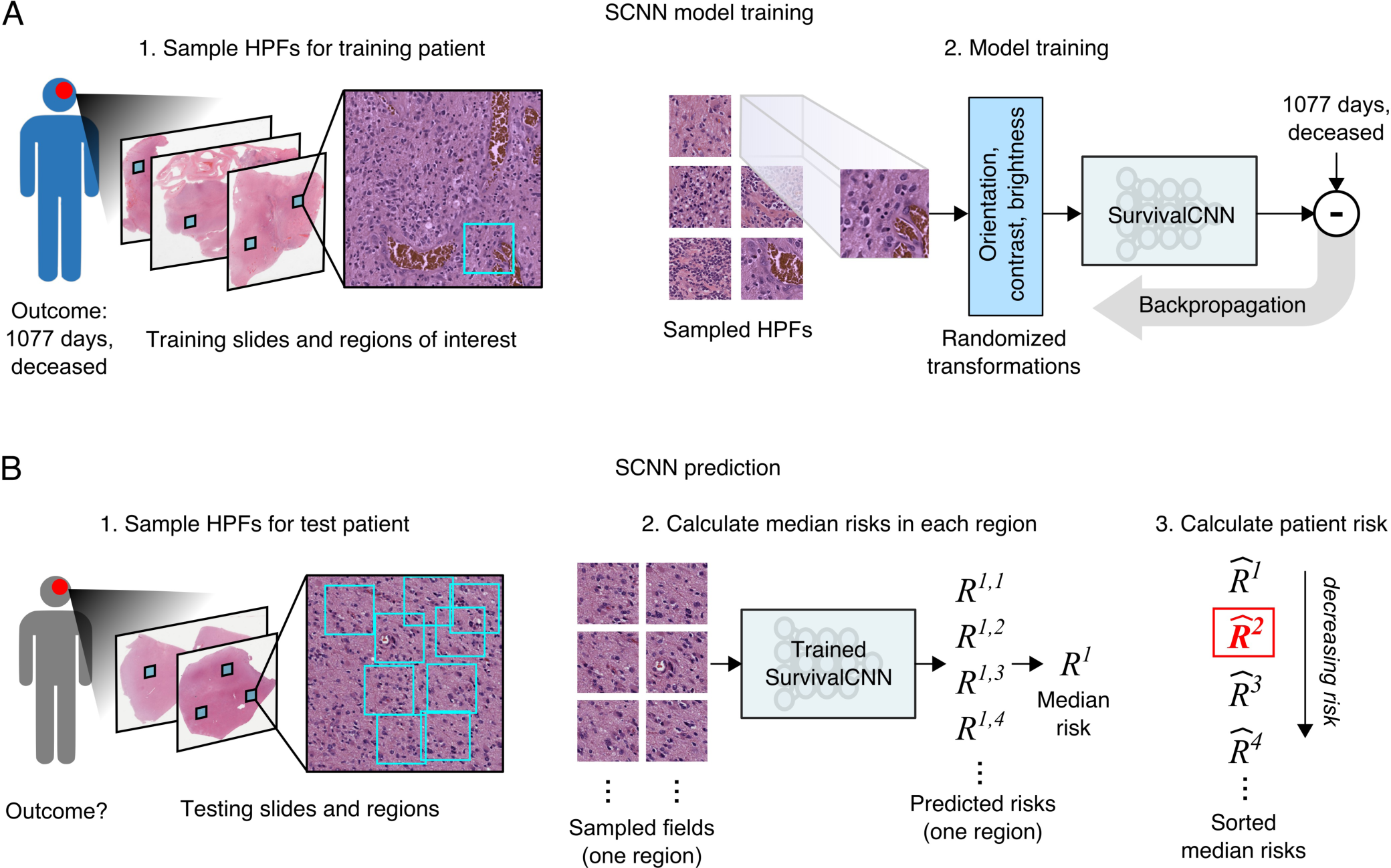
SCNN uses image re-sampling to improving the robustness of training and prediction. **(A)** During training, a single 256x256 pixel high-power field is sampled from each region, producing multiple HPFs per patient. Each HPF is subjected to a series of random transformations that simulate image acquisition variations, and is then used as an independent sample to update the network weights. New HPFs are re-sampled at each training epoch (one training pass through all patients). **(B)** When predicting the outcome of a newly diagnosed patient, 9 HPFs are sampled from each region of interest and a risk is predicted for each field. The median risk in each region is calculated, the median risks are sorted, and the second highest risk is selected as the risk of the patient. This process was designed to deal with tissue heterogeneity by emulating the process of histologic evaluation by a pathologist, where prognostication is based on the most malignant regions within a heterogeneous sample.

### Assessing the prognostic accuracy of SCNN

The prognostic accuracy of SCNN models was assessed using Monte Carlo cross validation. We first identified ROIs in 106 whole slide images of hematoxylin and eosin stained sections obtained from 769 gliomas Patients were assigned to either training (80%) validation (20%) to form 15 randomized datasets to evaluate the prognostic accuracy of methods (see Supplementary Tables S1 2). Accuracy was measured using Harrell’s c index, a non parametric statistic that measures concordance between predicted risks and actual survival (31) A c index of 1 indicates perfect concordance between predicted risk and overall survival, and a c index of 0.5 corresponds to random concordance

SCNN networks demonstrated substantial prognostic power, achieving a median c ndex of **0.754** (see **Figure 3B**) For comparison, we also measured the accuracy of baseline models generated using the genomic biomarkers and manual histologic grading used in the World Health Organization (WHO) classification (see **Figure 3A**) The WHO assigns gliomas to three genomic subtypes defined by mutations in isocitrate dehydrogenase (*IDH1* / *IDH2*) and co deletion of chromosomes 1p and 19q. Within these molecular subtypes, gliomas are further assigned a histologic grade based on criteria that vary depending on cell of origin (either astrocytic or oligodendroglial). These criteria include mitotic activity, nuclear atypia, the presence of necrosis, and the characteristics of microvascular structures. WHO baseline models based on molecular subtype and manual histologic grade had a median c index of **0.774** outperforming SCNN networks based on machine learning from histology images (Wilcoxon signed rank **p=2.61e-3**) The manual histologic grade baseline models had a median c index of **0.745** with performance similar to SCNN models (**p=0.307**). The molecular subtype baseline models had a median c index of **0.746**, and were significantly outperformed by the SCNN models (**p=4.68e-2**)

**Figure 3.**
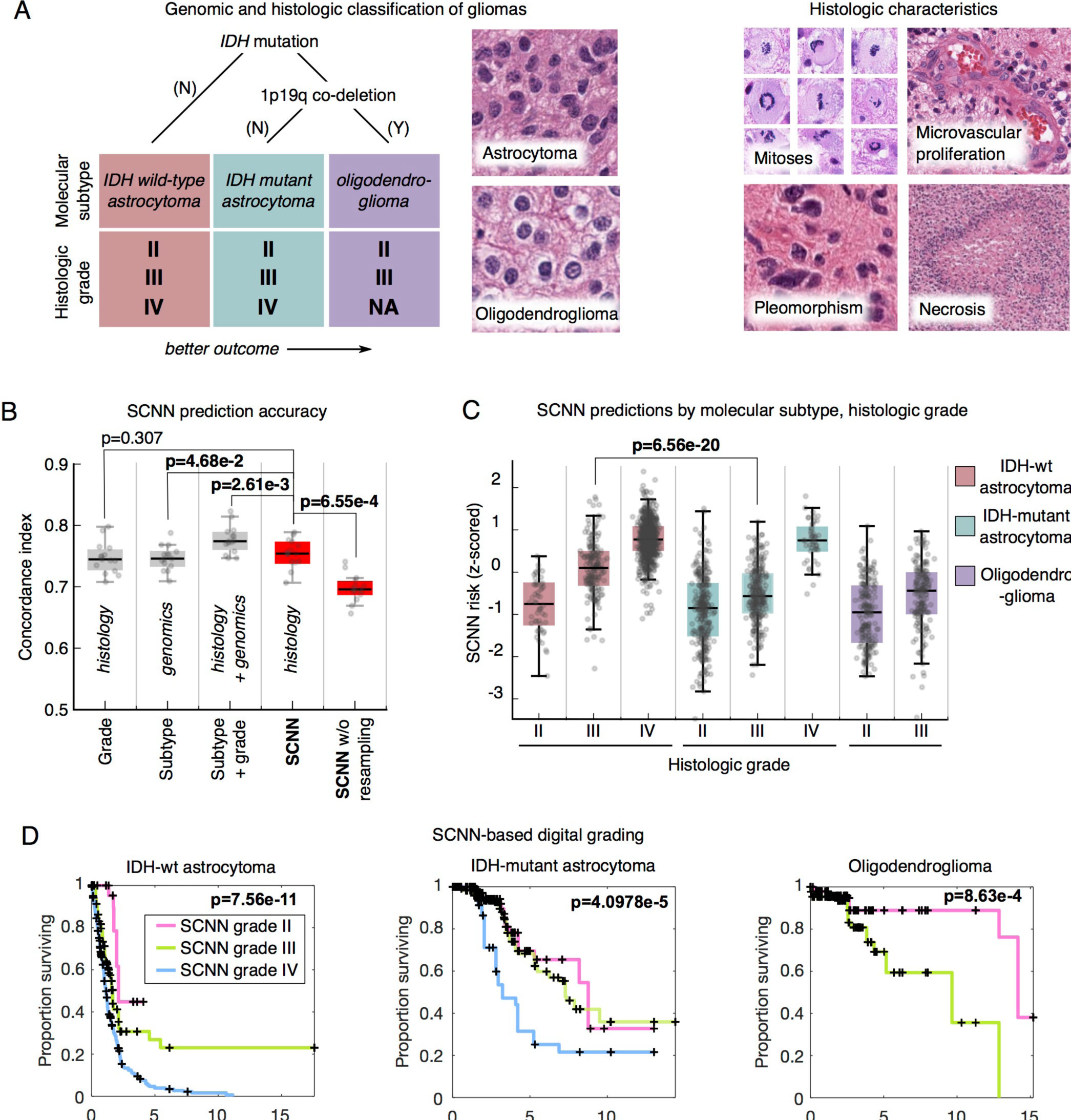
Prognostication criteria for diffuse gliomas. **(A)** Prognosis in the diffuse gliomas is determined by genomic classification and manual histologic grading. Tumors are first classified into three molecular subtypes based on the presence of mutations in *IDH1* / *IDH2* and the co-deletion of chromosomes 1p and 19q. Grade is then determined within each subtype using histologic characteristics. Subtypes with an astrocytic lineage are split by mutations in IDH, where co-deletion and IDH mutation defines oligodendroglial differentiation. These lineages associate with histologic patterns, however, histologic evaluation is not a reliable predictor of molecular subtype. Histologic criteria used for grading range from nuclear morphology to higher-level patterns like necrosis or the presence of abnormal microvascular structures. **(B)** Comparison of the prognostic accuracy of SCNN models to baseline models based on molecular subtype, and molecular subtype and histologic grade. Models were evaluated over 10 independent training/testing sets with randomized patient assignments, and with/without training and testing resampling. **(C)** The risks predicted by the SCNN models correlate with both histologic grade and molecular subtype, decreasing with grade and generally trending with the clinical aggressiveness of genomic subtypes. **(D)** Kaplan-Meier plots for digital grades based on risks predicted by SCNN models. SCNN models can stratify patients within each subtype similar to histologic grade (see **Figure S2**).

We also evaluated the benefits of our resampling methods in improving the performance of SCNN models Repeating the SCNN experiments without resampling techniques reduced the median c index to **0.696** significantly worse than for SCNN models where resampling was used (**p=6.55e −4**)

### SCNN predictions correlate with genomic subtypes and manual histologic grade

To further investigate the relationship between SCNN predictions and the WHO paradigm, visualized how risks predicted by SCNN networks are distributed across molecular subtype and histologic grade (see **Figure 3C**) SCNN predictions were highly correlated with both subtype and grade, and were consistent with expected patient outcomes in each category Firstly, within each molecular subtype, the risks predicted by SCNN increase with histologic grade Secondly, predicted risks are consistent with the published expected overall survivals associated with genomic subtypes (32) Astrocytomas with wild type IDH are highly aggressive with a median survival of 18 months, and the collective risks for these patients is higher than for patients from other subtypes. Astrocytomas having IDH mutations are another subtype with considerably better overall survival ranging from 3-8 years, and the predicted risks for patients in this subtype are more moderate Notably in this subtype, SCNN risks are not well separated for grades II and III consistent with reports of histologic grade being an inadequate predictor of outcome in this subtype (33) Gliomas with mutations in IDH and co deletion of chromosomes 1p/19q are described as *oligodendroglioma*, have a distinct differentiation, and have the lowest overall predicted risks consistent with survivals of 10+ years for this subtype Finally, we noted a significant difference in predicted risks for grade III gliomas in the astrocytic subtypes (rank sum **p=6.56e 20**). These subtypes share an astrocytic lineage are graded using identical histologic criteria, and are not known to have any distinguishing histological characteristics. Despite the inability of pathologists to discriminate between these subtypes using histology, SCNN can predict risks that are consistent with worse outcomes for grade III IDH wild type astrocytomas (median survival 1.7 years) compared to grade III IDH mutant astrocytomas (median survival 6.3 years).

To illustrate how SCNN risks can be used to assign a categorical “digital” grade, we performed a Kaplan Meier analysis to stratify patients based on SCNN risks (see **Figure 3D**). Risk thresholds defining digital grades were established for each molecular subtype separately. The proportions of each histologic grade in each subtype were used as a guideline to set thresholds on SCNN risks (see Methods) n each subtype, the digital grades capture survival differences in a manner analogous to manual histologic grading. A comparison to stratification by histologic grade is presented in **Supplementary Figure S2** Digital and manual histologic grades have similar prognostic power in IDH wild type astrocytomas (log rank **p=1.23e 12** versus **p=7.56e 11** respectively). In IDH mutant astrocytomas both digital and manual histologic grades have difficulty separating Kaplan Meier curves for grades II and III, yet both clearly distinguish grade IV as being associated with worse outcomes Discrimination for oligodendroglioma survival is also similar between digital and manual histologic grades (log rank **p=9.73e 7** versus **p=8.63e 4** respectively)

### Improving prognostic accuracy by integrating genomic biomarkers

To leverage both histologic and genomic data in predicting survival, we developed a hybrid Genomic SCNN model (GSCNN). The GSCNN learns prognosis from both genomics and histology by incorporating genomic variables into the fully connected layers of the SCNN to improve prognostic accuracy (see **Figure 4**). This configuration enables the genomic variables to influence the patterns learned from histology by providing information on molecular subtype near the terminal network layers.

**Figure 4.**
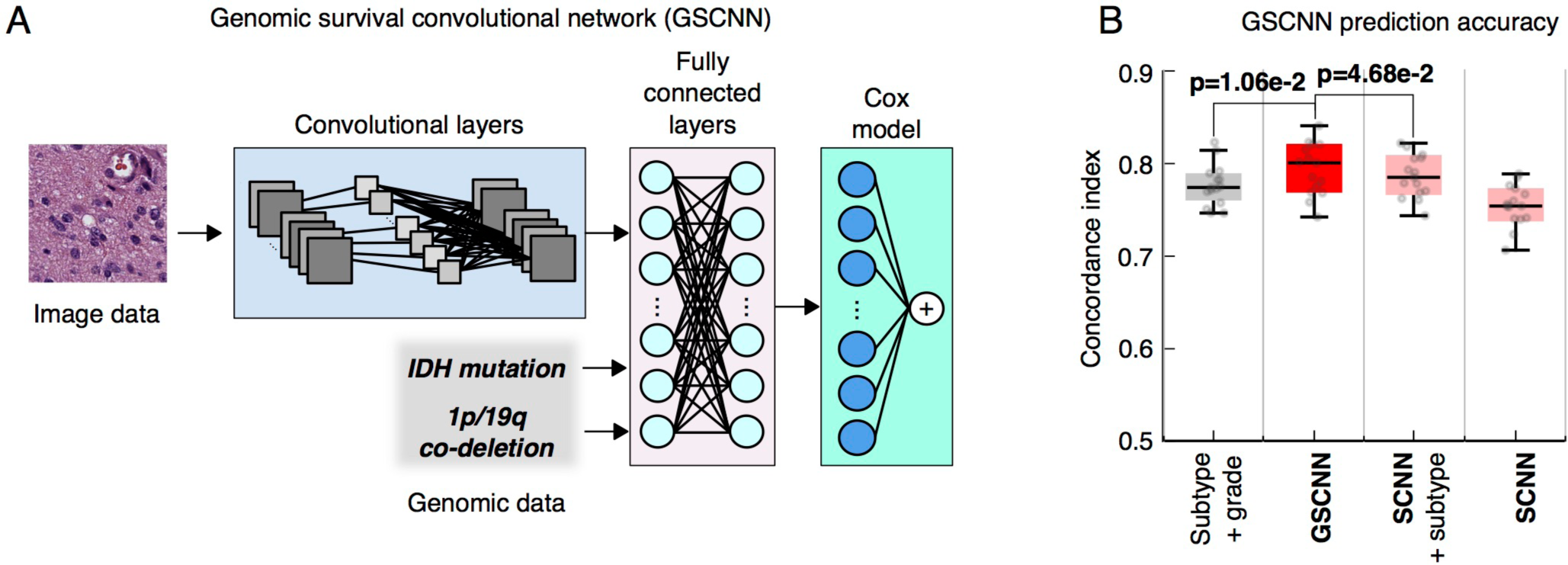
Genomic-SCNN models integrate genomic and imaging data for improved performance. **(A)** A hybrid architecture was developed to combine histology image and genomic data to make integrated predictions of patient survival. These models incorporate genomic variables as inputs to their fully-connected layers. Here, we show the incorporation of genomic variables for gliomas, however any number of genomic or proteomic measurements can be similarly used. **(B)** The GSCNN models significantly outperform SCNN models, as well as the WHO paradigm based on genomic subtype and histologic grading.

We repeated our experiments using GSCNN models with histology images, IDH mutation status, and 1p/19q co-deletion as inputs, and found that the median c-index improved to **0.801**. The addition of genomic variables improved the performance by 5% on average over SCNN models that are trained on histology images alone. The GSCNN models also significantly outperform the WHO baseline subtype-grade model trained on equivalent data (signed-rank **p=1.06e-2**). We compared the GSCNN approach to a more superficial way of integrating genomic variables, where a Cox model was trained using SCNN risks and the IDH and 1p/19q variables. This superficial approach did not perform as well as GSCNN, with a median c-index of **0.785** (**p=4.68e-2**).

To evaluate the independent prognostic power of risks predicted by SCNN and GSCNN, we performed a multivariable Cox regression analysis (see **Table 1**). In a multivariable regression that included SCNN risks, subtype, grade, age, and sex, SCNN risks were prognostic when correcting for all other features including manual grade and molecular subtype (**p=2.71e-12**). Manual histologic grade was not significant in this regression analysis. We also performed a similar multivariable regression with GSCNN risks, and found GSCNN to be significant (**p=9.69e-12**). In this multivariable regression, molecular subtype was not significant, and histologic grade was only marginally significant.

**Table 1.** Hazard ratios for univariable and multivariable Cox regression models.

| Variable | Single variable |  |  |  | Multivariable (SCNN) |  |  | Multivariable (GSCNN) |  |  |
| --- | --- | --- | --- | --- | --- | --- | --- | --- | --- | --- |
|  | C-index | Hazard Ratio | 95% CI | p-value | Hazard Ratio | 95% CI | p-value | Hazard Ratio | 95% CI | p-value |
| **SCNN** | 0.741 | 7.15 | (5.64, 9.07) | **2.08e-61** | 3.05 | (2.22, 4.19) | **2.71e-12** | - | - | - |
| **GSCNN** | 0.781 | 12.60 | (9.34, 17.0) | **3.08e-64** | - | - | - | 8.83 | (4.66, 16.74) | **9.69e-12** |
| **IDHwt astrocytoma** | 0.72556 | 9.21 | (6.88, 12.34) | **3.48e-52** | 4.73 | (2.57, 8.70) | **3.49e-7** | 0.97 | (0.43, 2.17) | 0.93 |
| **IDHmut astrocytoma** | - | 0.23 | (0.170, 0.324) | **2.70e-19** | 2.35 | (1.27, 4.34) | **5.36e-3** | 1.67 | (0.90, 3.12) | 0.10 |
| **Histologic Grade IV** | 0.72106 | 7.25 | (5.58, 9.43) | **2.68e-51** | 1.52 | (0.839, 2.743) | 0.159 | 1.98 | (1.11, 3.51) | **0.017** |
| **Histologic Grade III** | - | 0.44 | (0.332, 0.591) | **1.66e-08** | 1.57 | (0.934, 2.638) | 0.0820 | 1.78 | (1.07, 2.97) | **0.024** |
| **Age** | 0.74413 | 1.77 | (1.63, 1.93) | **2.52e-42** | 1.33 | (1.20, 1.47) | **9.57e-9** | 1.34 | (1.22, 1.48) | **9.30e-10** |
| **Sex (F)** | 0.55162 | 0.89 | (0.706, 1.112) | 0.29 | 0.85 | (0.67, 1.08) | 0.168 | 0.86 | (0.68, 1.08) | 0.18 |

### Visualizing prognosis with SCNN heatmaps

Deep learning networks are often criticized for being “black-box” approaches that do not reveal insights into their prediction mechanisms. To investigate the visual patterns SCNN models learn from histology images, we created *risk heatmap* overlays to visualize the risks associated with different regions in a whole-slide image. These heatmaps were generated by predicting risk for each non-overlapping HPF in a whole-slide image. The predicted risks of each HPF were used to generate a color-coded transparent overlay that represents the SCNN risk predictions across the entire slide.

A selection of risk heatmaps from three patients is presented in Figure 5, with inlays demonstrating how SCNNs associate risk with important pathologic phenomena. For TCGA-DB-5273 (Grade III, IDH mutant astrocytoma), the SCNN heatmap clearly highlights regions of early microvascular proliferation, an advanced form of angiogenesis that is a hallmark of malignant progression, as being associated with high risk. Risk in this heatmap also increases with cellularity, heterogeneity in nuclear shape and size (pleomorphism), and the presence of abnormal microvascular structures. Regions in TCGA S9 A7J0 have varying extents of tumor infiltration, ranging from normal brain, to sparsely infiltrated adjacent normal regions exhibiting satellitosis (where neoplastic cells cluster around neurons), to moderately and highly infiltrated regions. This heatmap correctly associates the lowest risks to normal brain regions, and can distinguish normal brain from adjacent regions that are sparsely infiltrated. Interestingly, higher risks are assigned to sparsely infiltrated regions (region 1, upper panel) than to regions with containing relatively more tumor infiltration (region 2). We observed a similar pattern in TCGA-TM-A84G, where edematous regions (region 1, lower panel) adjacent to moderately cellular tumor regions (region 1, upper panel) are also assigned higher risks. These latter examples provide novel risk features embedded within histologic sections that have been previously unrecognized and could inform and improve pathology practice.

**Figure 5.**
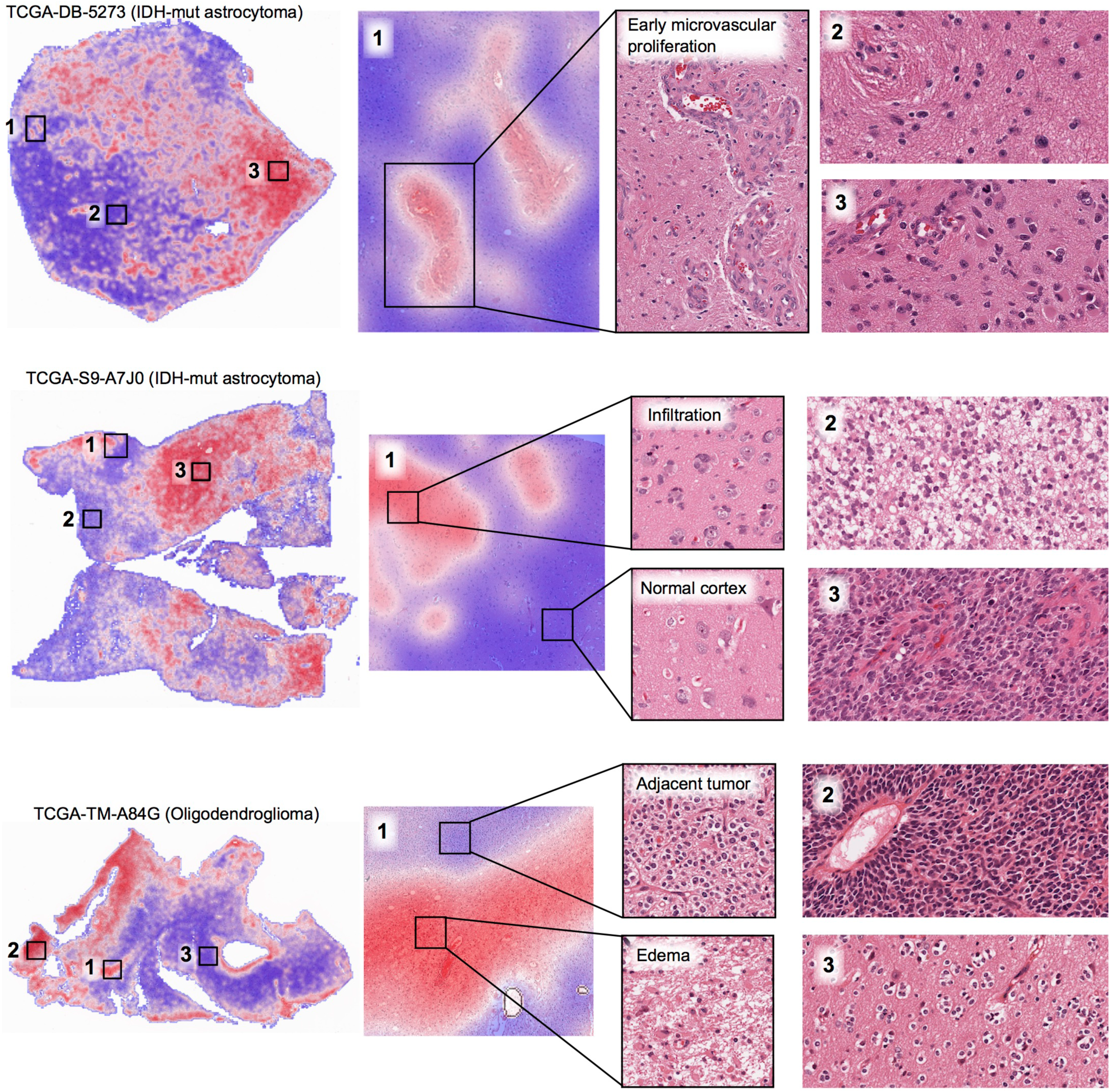
Visualizing risk with whole-slide SCNN heatmaps. We performed SCNN predictions exhaustively within whole slide images to generate heatmap overlays of the risks that SCNN associates with different histologic patterns. Red indicates relatively higher risk, and blue lower risk (the scale for each slide is different). (Top) In TCGA-DB-5273, SCNN clearly and specifically associates early microvascular proliferation with high-risks (region 1), and also higher risks with increasing tumor infiltration and cell density (region 2 versus 3). (Middle) In TCGA-S9-A7J0, SCNN can appropriately discriminate between normal cortex (region 1, lower panel) and adjacent regions infiltrated by tumor (region 1, upper panel). Highly cellular regions containing prominent microvascular structures (region 3) are again assigned higher risks than lower density regions of tumor (region 2). Interestingly, low density infiltrate in the cortex was associated with high risk (region 1, upper panel) (Bottom) In TCGA-TM-A84G, SCNN assigns high risks to edematous regions (region 1, lower panel) that are adjacent to tumor (region 1, upper panel).

## DISCUSSION

We developed a deep learning algorithm for learning survival directly from histological images, and systematically evaluate its prognostic accuracy in the context of the current clinical standard based on genomic classification and histologic grading. In contrast to a previous study that achieved only very marginal prediction accuracy, SCNN rivals or exceeds the performance of highly trained human experts in assessing prognosis.

Our study provides new insights into applications of artificial intelligence in medicine, and also new technical approaches for dealing with intratumoral heterogeneity and training data deficits. We also developed a visualization technique that allows pathologists to explore the associations between histological patterns and prognosis over large whole slide images that inevitably exhibit significant heterogeneity.

Our study investigated the ability to predict overall survival in gliomas, a disease with wide variations in outcomes, and an ideal test case where histologic grading and genomic classifications have independent prognostic power. Remarkably, SCNN performed as well as manual histologic grading or molecular subtyping in predicting overall survival in our dataset. Further investigation of the associations between SCNN risk predictions, genomic subtypes, and histologic grades revealed that SCNN can effectively discriminate outcomes in each subtype, effectively performing digital histologic grading. Furthermore, SCNN can effectively recognize differences in images that associate with genomic subtypes, and predict risks accordingly. Oligodendrogliomas have a distinct histology, and so the ability to discriminate this subtype is not unexpected. For astrocytomas, the SCNN network could correctly predict higher risks for grade III IDH wild-type astrocytomas than for grade III IDH mutant astrocytomas, suggesting that SCNN may be able to recognize subtle histologic differences associated with IDH mutations that are not yet appreciated by pathologists. The broad hypermethylation induced by IDH mutations could plausibly affect nuclear appearance, providing a possible explanation of visual differences that are detectable by SCNN, but more investigation of this topic is needed.

To integrate genomic information in prognostication, we developed a hybrid approach that learns from both histology images and genomic biomarkers. The GSCNN model presented in our study *significantly outperforms the WHO standard based on identical inputs*. By providing molecular subtype data directly to the network, instead of relying on inferences from histology, GSCNN can focus more attention on learning histologic patterns associated with disease progression in each subtype. This result illustrates how complementary genomic and image data can be practically integrated into a single prediction framework, an issue that presents a significant barrier in the clinical implementation of computational prognostication. Our previous work in developing deep-learning survival models from genomic data has shown that accurate survival predictions can be learned from high-dimensional genomic and protein expression signatures (34). Incorporating additional genomic variables into GSCNN models is an area for future research, and requires larger data sets with both histologic images and rich genomic annotations.

While deep learning methods frequently deliver outstanding performance, the interpretability of these models is limited, and remains a significant barrier in their validation and adoption. The risk heatmap provides insights into the histologic patterns associated with increased risk, and can also serve as a practical tool to guide pathologists to tissue regions associated with worse prognosis. This approach suggests that our network can learn visual patterns associated with histologic criteria used in grading including microvascular proliferation, cell density, and nuclear morphology. Microvascular prominence and proliferation are associated with disease progression in all forms of diffuse glioma, and these features are clearly delineated as high-risk in the heatmap presented for TCGA-DB-5273. Likewise, increases in cell density and nuclear pleomorphism are also associated with increased risk in all examples. In addition to these results, the heatmap analysis provided some interesting results that need to be further investigated. In region 1 of TCGA-S9-A7J0, SCNN assigns higher risk to sparsely infiltrated cerebral cortex than to region 2 that is infiltrated by a higher density of tumor cells (adjacent normal cortex in region 1 is properly assigned a very low risk). Widespread infiltration into distant sites of the brain is a hallmark of gliomas, and results in treatment failure since surgical resection of visible tumor leaves residual neoplastic infiltrates. It is not clear that this is the reason for SCNN assigning high risk to sparsely infiltrated regions, but nonetheless, it is an interesting finding worth pursuing. Similarly, region 1 of TCGA-TM-A84G illustrates a high risk associated with low cellularity edematous regions, compared to adjacent oligodendroglioma with much higher cellularity. Edema is frequently observed within gliomas and in adjacent brain and its degree may be related to the rate of growth (35) yet its histologic presence has not been previously recognized as a feature of aggressive behavior or incorporated into grading schemes. These observations confirm that risks predicted by SCNN are not purely a function of cellular density or nuclear atypia and demonstrate that these methods can identify novel, potentially practice changing features associated with increased risk embedded within pathology images.

Although our study provides insights into deep learning for precision medicine, it has some important limitations. A relatively small portion of each slide was used for training, and the selection of regions of interest requires expert guidance. More advanced methods are needed for automatically selecting regions and for incorporating more of the slide into the learning and prediction process. A single whole slide image can contain remarkable heterogeneity, and so incorporating more of the slide into the learning process will require more advanced training methods. Our method currently produces a dimensionless risk by optimizing partial likelihood, and learning of the baseline hazard would permit calibrated prediction of actual survival times. Finally, while we have applied our techniques to gliomas, validation of these approaches in other diseases is needed and could provide additional insights. Furthermore, our methods are not specific to histology imaging or cancer applications, and could be adapted to other medical imaging modalities and biomedical applications.

## METHODS

### Data and image curation

Whole slide images, clinical and genomic data were obtained from The Cancer Genome Atlas via the Genomic Data Commons (https://gdc.cancer.gov/). Images of diagnostic hematoxylin and eosin stained formalin-fixed paraffin-embedded sections from the Brain Lower Grade Glioma (LGG) and the Glioblastoma (GBM) cohorts were reviewed to remove images containing tissue processing artifacts including bubbles, section folds, pen markings and poor staining. Representative regions of interest containing primarily tumor nuclei were manually identified for each slide that passed quality control. In the case of grade IV disease, some regions include microvascular proliferation as this feature was exhibited throughout tumor regions. Regions containing geographic necrosis were excluded. A total of 1061 whole-slide images from 769 unique patients were analyzed.

Regions of interest images (1024x1024 pixels) were cropped at 20X objective magnification using OpenSlide and color-normalized to a gold-standard H&E calibration image to improve consistency of color characteristics across slides. High-power fields (HPFs) at 256x256 pixels were sampled from these regions and used for training and testing as described below.

### Network architecture and training procedures

The survival convolutional neural network combines elements of the 19-layer VGG convolutional network architecture with a Cox proportional hazard model to predict time-to-event data from images (see **Figure S1**) (36). Image feature extraction is achieved by four groups of convolutional layers: 1) The first group contains two convolutional layers with 64 3x3 kernels, interleaved with local normalization layers, then followed with a single max-pooling layer 2) The second group contains two convolutional layers (128 3x3 kernels) interleaved with two local normalization layers, followed by a single max pooling layer 3) The third group interleaves four convolutional layers (256 3x3 kernels) with four local normalization layers, followed by a single max pooling layer 4) The fourth group contains interleaves of eight convolutional (512 3x3 kernels) and eight local normalization layers, with an intermediate and a terminal max pooling layer. These four groups are followed by a sequence of 3 fully connected layers containing 1000, 1000, and 256 nodes respectively.

The terminal fully connected layer outputs a prediction of risk *R* = β^*T*^ *X* associated with the input image, where β ∈ℝ^256 × 1^ - are the terminal layer weights and % ∈ℝ^256 × 1^ are the inputs to this layer. To provide an error signal for backpropagation, these risks are input to a Cox proportional hazards layer to calculate the negative partial log-likelihood

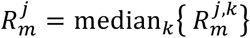

where *β*^*T*^%_*i*_is the risk associated with HPF *i*, *U* is the set of right-censored samples, and Ω_*i*_is the set of “at-risk” samples with event or follow-up times Ω_i_= {*j* | *Y*_*j*_:≥ *Y*_*i*_(where *Y*_*i*_ is the event or last follow-up time of patient *i*).

The adagrad algorithm was used to minimize the negative partial log-likelihood via backpropagation to optimize model weights, biases and convolutional kernels (37). Parameters to adagrad include the initial accumulator value = 0.1, initial learning rate = 0.001, and an exponential learning rate decay factor = 0.1. Model weights were initialized using the variance scaling method (38), and a weight decay was applied to the fully connected layers during training (decay rate = 4e-4). Models were trained for 100 epochs (1 epoch is one complete cycle through all training samples) using mini-batches consisting of 14 HPFs each. Each mini-batch produces a model update, resulting in multiple updates per epoch. Calculation of the Cox partial likelihood requires access to the predicted risks of all samples, which are not available within any single mini-batch, and so Cox likelihood was calculated locally within each mini-batch to perform updates (*U* and Ω_*i*_were restricted to samples within each mini-batch). Local likelihood calculation can be very sensitive to how samples are assigned to mini-batches, and so we randomize the mini-batch sample assignments at the beginning of each epoch to improve robustness. Mild regularization was applied during training by randomly dropping out 5% of weights in the last fully connected layer in each mini-batch during training to mitigate overfitting.

### Training resampling

Each patient has possibly multiple slides, and multiple regions within each slide that can be used to sample HPFs. During training, a single HPF was sampled from each region, and these HPFs were treated as semi-independent training samples. Each HPF was paired with patient outcome for training, duplicating outcomes for patients containing multiple regions / HPFs. The HPFs are resampled at the beginning of each training epoch to generate an entirely new set of HPFs. Randomized transforms were also applied to these HPFs to improve robustness to tissue orientation and color variations. Since the visual patterns in tissues can often be anisotropic, we randomly apply a mirror transform to each HPF. We also generate random transformations of contrast and brightness using the “random_contrast” and “random_brightness” TensorFlow transformations to modify the HPF and simulate color variations. These resampling and transformation procedures, along with the use of multiple HPFs for each patient, has the effect of augmenting the effective size of the labeled training data. Similar approaches for training data augmentation have demonstrated considerable improvements in general imaging applications (39).

### Testing resampling and model averaging

Resampling was also performed to increase the robustness and stability of predictions: 1) 9 high-power fields are first sampled from each region *j* corresponding to patient E 2) The risk of the k^th^ HPF in region *j* for patient *m*, denoted 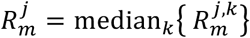, is then calculated using the trained SCNN model 3) The median risk 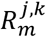 is calculated for region *j* using the aforementioned HPFs to reject outlying risks 4) These median risks are then sorted from highest to lowest 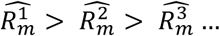, where the superscript index now corresponds to the risk rank 5) The risk prediction for patient *m* is then selected as the second highest risk 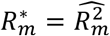 This filtering procedure was designed to emulate how a pathologist integrates information from multiple areas within a slide, determining prognosis based on the region associated with the worst prognosis. Selection of the second highest risk (as opposed to the highest risk) introduces robustness to outliers or high risks that may occur due to some imaging or tissue-processing artifact.

Since the accuracy of our models can vary significantly from one epoch to another, largely due to the training resampling and randomized mini-batch assignments, a model averaging technique was used to reduce prediction variance. To obtain final risk predictions for the testing patients that are stable, we perform model averaging using the models from epochs 96-100 to smooth variations across epochs and increase stability. Formally, the model-averaged risk for patient *m* is calculated as

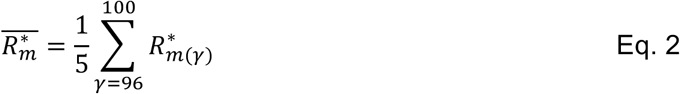

where 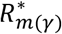 denotes the predicted risk for patient E in training epoch *γ*.

### Validation procedures

Patients were randomly assigned to non-overlapping training (80%) and validation (20%) sets that were used to train models and evaluate their performance. If a patient was assigned to training, then all slides corresponding to that patient were assigned to the training set and likewise for the testing set. This ensures that no data from any one patient is represented in both training and testing sets to avoid overfitting and optimistic estimates of generalization accuracy. We repeated the randomized assignment of patients training/testing sets 15 times, and used each of these training/testing sets to train and evaluate a model. The same training/testing assignments were used in each model (SCNN, GSCNN, baseline) for comparability. Prediction accuracy was measured using Harrell’s c-index (CI) to measure the concordance between predicted risk and actual survival for testing samples (31).

### Statistical analyses

C-indexes generated by Monte Carlo cross-validation were performed using the Wilcoxon signed rank test. This paired test was chosen because each method was evaluated using identical training/testing sets. Comparisons of SCNN risk values across grade were performed using the Wilcoxon rank-sum test. Cox univariable and multivariable regression analyses were performed using predicted SCNN risk values for all training and validation samples in the randomized training/validation set 1. Analysis of the correlation of grade, molecular subtype, and SCNN risk predictions were performed by pooling predicted risks for validation samples across all experiments. SCNN risks were normalized within each experiment by z-score prior to pooling. Grade analysis was performed by determining “digital” grade thresholds for SCNN risks in each subtype. Thresholds were objectively selected to match the proportions of samples in each histologic grade in each subtype. Statistical analysis of Kaplan Meier plots was performed using the log-rank test.

### Hardware and software

Prediction models were trained using TensorFlow (v0.12.0) on servers equipped with dual Intel(R) Xeon(R) CPU E5-2630L v2 @ 2.40GHz CPUs, 128GB RAM, and dual NVIDIA K80 graphics cards. Image data was extracted from Aperio. svs whole-slide image formats using OpenSlide (http://openslide.org/). Basic image analysis operations were performed using HistomicsTK (https://github.com/DigitalSlideArchive/HistomicsTK), a Python package for histology image analysis.

### Data availability

This paper was produced using large volumes of publicly available genomic and imaging data. The authors have made every effort to make available links to these resources as well as making publicly available the software methods and information used to produce the datasets, analyses, and summary information. All data not published in the tables and supplements of this article are available from the corresponding author on request.

## AUTHOR CONTRIBUTIONS

PM and SY developed the primary method. PM, DAG, and JEVV curated datasets. PM, MA, LADC and DJB designed experiments. PM performed experiments. JBS performed statistical analyses and assisted with manuscript writing and editing. DJB assisted with the interpretation of results and manuscript writing. LADC conceived of the ideas and oversaw the work.

## ACKNOWLEDGEMENTS

This work was supported by the U.S. National Institutes of Health, National Library of Medicine Career Development Award K22LM011576, and National Cancer Institute grant U24CA194362, and the National Brain Tumor Society.

## SUPPLEMENTARY FIGURES

**Figure S1.**
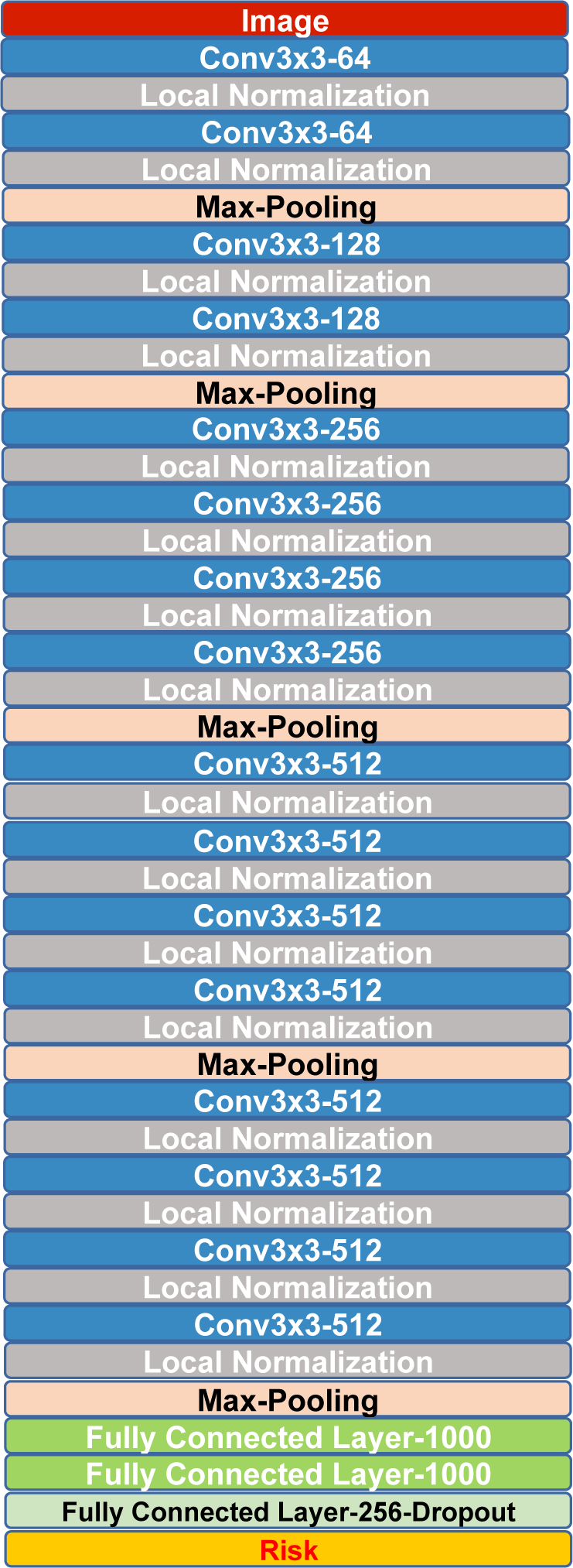
Detailed diagram of the survival convolutional neural network architecture. The architecture is a variation of the VGG network, and combines convolutional, max pooling, local normalization, and fully connected layers.

**Figure S2.**
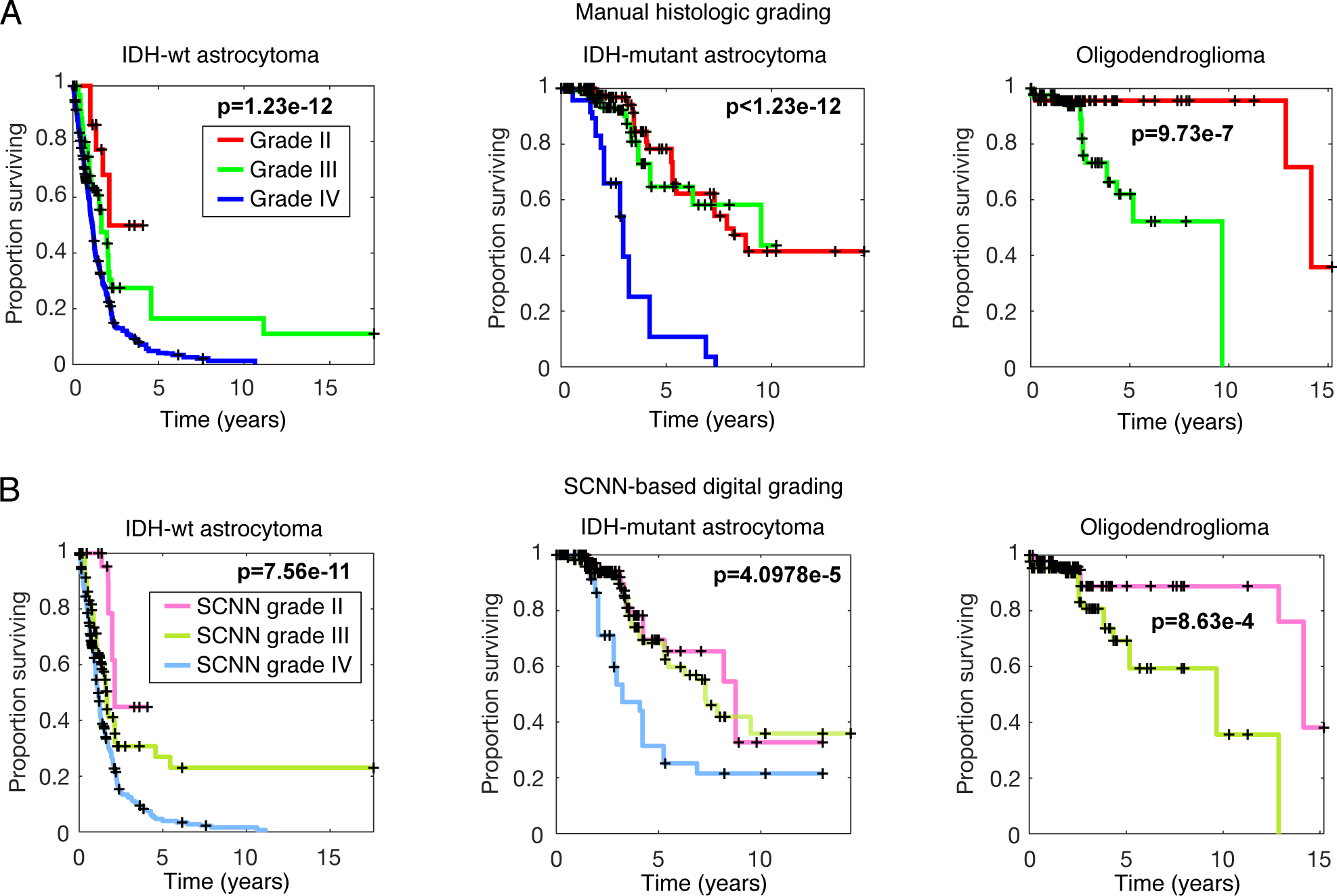
Comparison of manual histologic grading and “digital” grading. **(A)** Kaplan-Meier plots for manual histologic grading by a neuropathologist. One plot is provided for each genomic subtype. **(B)** Kaplan-Meier plots for digital grades based on risks predicted by SCNN models. Thresholds for digital grades were established using the proportion of samples in each histologic grade for each subtype. Histologic grading and grading by SCNN produce almost identical stratifications of patient survival. Notably, digital grading cannot discriminate survival between grades II/III in the IDH-mutant astrocytomas, just as manual histologic grading cannot.

